# Pervasive and Dynamic Transcription Initiation in *Saccharomyces cerevisiae*

**DOI:** 10.1101/450429

**Authors:** Zhaolian Lu, Zhenguo Lin

## Abstract

Transcription initiation is finely regulated to ensure the proper expression and function of these genes. The regulated transcription initiation in response to various environmental cues in the model organism *Saccharomyces cerevisiae* has not been systematically investigated. In this study, we generated quantitative maps of transcription start site (TSS) at a single-nucleotide resolution for *S. cerevisiae* grown in nine different conditions using no-amplification non-tagging Cap analysis of gene expression (nAnT-iCAGE) sequencing. Based on 337 million uniquely mapped CAGE tags, we mapped ~1 million well-supported TSSs, suggesting highly pervasive transcription initiation in the compact genome of yeast. The comprehensive TSS maps allowed us to identify core promoters for ~96% verified protein-coding genes and to revise the predicted translation start codon for 183 genes. We found that 56% of yeast genes have at least two core promoters and alternative usage of different core promoters in a gene is widespread in response to changing environments. More importantly, most core promoter shifts are coupled with differential gene expression, indicating that core promoter shift might play an important role in controlling transcriptional activity of yeast genes. Based on their dynamic activities, we divided yeast core promoters as constitutive core promoters (55%) and inducible core promoters (45%). The two classes of core promoters exhibit distinctive patterns in transcriptional abundance, chromatin structure, promoter shape, and sequence context. In summary, the quantitative TSS maps generated by this study improved the annotation of yeast genome, and revealed a highly pervasive and dynamic nature of transcription initiation in yeast.

## INTRODUCTION

The RNA polymerase II (pol-II) core promoter is the region where pol-II is recruited to initiate transcription. It includes the transcription start sites (TSSs) and immediately flanking sequences that contain various DNA motifs to accurately direct transcription initiation by the pre-initiation complex (PIC) (Butler and Kadonaga 2002). Core promoter is the final target of actions of almost all the factors involved in the transcriptional regulation because the regulatory signals of transcription are ultimately integrated to the initiation process at core promoters (Juven-Gershon and Kadonaga 2010). Accurate transcription initiation is vital to ensure the proper expression and function of these genes (Smale and Kadonaga 2003). Mis-regulation of transcription initiation has been found to be associated with a broad range of human diseases, such as breast cancer, diabetes, kidney failure and Alzheimer’s disease (Arrick et al. 1991; Romeo et al. 1993; Capoulade et al. 2001; Sobczak and Krzyzosiak 2002; Mihailovich et al. 2007). In this regard, accurate identification of TSSs and characterization of their regulated activities are essential for obtaining fundamental insights into regulatory mechanisms that determine the location and activities of transcription initiation. The global maps of TSS and core promoter have been generated in several important metazoans, such as human (Consortium 2012), mouse (Carninci et al. 2005), fruit fly (Hoskins et al. 2011) and zebrafish (Haberle et al. 2015). These maps revealed that most animal genes contain multiple core promoters and the selections and activities of core promoters are precisely regulated to ensure that a correct transcript is produced at an appropriate level in a tissue or developmental stage (Carninci et al. 2006; FANTOM Consortium and the RIKEN PMI and CLST (DGT) et al. 2014).

The budding yeast, *Saccharomyces cerevisiae* has served as eukaryotic model organisms for many landmark discoveries in gene regulation mechanisms and other cellular processes over the past several decades (Duina et al. 2014). Various techniques have been applied to identify genome-wide TSS for *S. cerevisiae*, such as microarray (Hurowitz and Brown 2003; David et al. 2006; Xu et al. 2009), SAGE (serial analysis of gene expression) (Zhang and Dietrich 2005), sequencing of full-length cDNA clones (Miura et al. 2006), RNA sequencing (Nagalakshmi et al. 2008; Waern and Snyder 2013), Transcript isoform sequencing (TIF-seq) (Pelechano et al. 2013), transcript-leaders sequencing (TL-seq) (Arribere and Gilbert 2013), and modified 5’RACE (Malabat et al. 2015). However, regarding generating accurate, comprehensive and quantitative TSS maps, most of these studies have limitations in resolutions, depth of coverage or sensitivity (Boyle et al. 2004; Grabherr et al. 2011; Batut et al. 2013; Steijger et al. 2013; Wade and Grainger 2014). For instance, microarray is known to have limited resolution and sensitivity for identifying 5’ boundary of transcripts (Wade and Grainger 2014). SAGE and full-length cDNA sequencing lack the throughput to provide sufficient data for lowly expressed genes and the quantitative measurements of TSS usage (Shiraki et al. 2003).

Tuning gene expression is an essential way to maximize cell survival through rapid responses to environmental stresses, particularly for unicellular organisms (de Nadal et al. 2011). Therefore, studying the activity of core promoters in response to changing environments is important for gaining fundamental insights into regulatory mechanisms of transcription initiation in yeast. Previous studies based on TIF-seq, TL-seq, and modified 5’RACE, were performed in only one or two growth conditions (Arribere and Gilbert 2013; Pelechano et al. 2014; Malabat et al. 2015). Consequently, the TSSs and core promoters identified in these studies may only represent a small part of transcription initiation landscape in yeast, as many transcripts being generated only in particular growth conditions. It is also challenging to accurately characterize the dynamic activities of core promoters with one or two examined conditions. In one study, RNA-sequencing has been carried out in *S. cerevisiae* grown in 18 different conditions and extensive different 5’ends have been observed, suggesting the dynamic of transcription initiation in yeast under different growth conditions (Waern and Snyder 2013). However, RNA-seq is known to have a shortcoming of inaccurate determination of TSSs, as assembly of RNA-seq reads usually extends transcript contigs to the very 5’ end, which lack the information of other TSSs within the longest transcript (Batut et al. 2013; Steijger et al. 2013; Boley et al. 2014; Wade and Grainger 2014). Therefore, it is necessary to generate high-resolution and quantitative TSS maps for yeast cells grown under various conditions to better understand the regulated dynamic of transcription initiation.

A revised Cap analysis gene expression (CAGE) technique, called no-amplification non-tagging CAGE libraries for Illumina sequencers (nAnT-iCAGE), is ideal for generating TSS maps at a single-nucleotide resolution and simultaneously quantifying their activities (Murata et al. 2014). Similar to TIF-seq (Pelechano et al. 2013) and modified 5’RACE (Malabat et al. 2015), nAnT-iCAGE captures the 7-methylguanosine cap structure at the 5’end of transcripts, and sequence the transcripts using high-throughput sequencers. By mapping sequenced reads to a reference genome, the exact TSS locations can be identified. The number of reads mapped to a TSS also quantifies the number of transcripts initiated at the TSS. Moreover, nAnT-iCAGE does not involve PCR amplification or restriction enzyme digestion (Murata et al. 2014), which are required for TIF-seq. Amplification by PCR has sequence-dependence efficiency which might introduce bias on transcription level. Restriction enzyme digestion may cause loss of RNA samples. Another limitation of TIF-seq is its bias toward short RNA molecules, which is also a common problem of full-length cDNA sequencing approach (Miura et al. 2006).

With a goal to characterize the regulatory dynamic of transcription initiation with an unprecedented resolution and depth, we used the nAnT-iCAGE technique to generate quantitative TSS maps for *S. cerevisiae* grown under nine different conditions that simulate their natural environments. Sampling transcripts from different growth conditions increase the number of identified TSSs or core promoters, providing a more complete picture of transcription initiation landscape in the unicellular model organism. The high-resolution TSS maps also allowed us to revise incorrected predicted translation start codon for many open reading frames (ORFs), improving the accuracy of genome annotation. Furthermore, our comparative analysis of TSS maps revealed the activities of different core promoters in a gene are highly dynamic in response to changing environments. Based on their transcriptional activities, we identified two distinct types of core promoters: the constitutive core promoters and inducible core promoters. The two types of core promoters differ from each other in transcriptional abundance, chromatin structure, sequence context, suggesting the presence of different regulatory strategies of transcription initiation in eukaryotic organisms.

## RESULTS

### Pervasive transcription initiations in *S. cerevisiae*

The *S. cerevisiae* strain BY4741, a haploid derivative of laboratory strain S288c, was used as the study system to generate high-resolution TSS maps. The 5’boundary of transcripts was captured following the nAnT-iCAGE protocol for *S. cerevisiae* cells grown in nine conditions (Table 1), which are informative on the natural environments and common stresses of wild yeast populations. For each growth condition, two biological replicates of nAnT-iCAGE libraries were constructed (18 libraries in total). All nAnT-iCAGE libraries were sequenced using Illumina NextSeq 500 (single-end, 75-bp reads), which yielded 636 million sequencing tags in total (Table S1), providing an unprecedented depth of coverage for 5’boundary of transcripts in yeast.

With a mapping rate of 91.9%, 584,689,028 tags were aligned to the reference genome of *S. cerevisiae* (assembly R64-2-1). We only used tags that are uniquely mapped to the reference genome (337,237,124 tags) for further analysis. The Pearson correlation coefficient *r* of the tag counts of the CTSSs (CAGE tags identified TSS) between the two biological replicates of each growth condition range from 0.97 to 1 (Figure S1), supporting the high reproducibility of the nAnT-iCAGE technique. Systematic G nucleotide addition bias at the 5’end of CAGE tags was corrected based on the probability of G addition (Carninci et al. 2006). The numbers of CTSSs identified in each growth condition range from 1,106,287 to 1,632,079 (Table S2). A total number of unique CTSSs is 4,254,561 after combining data from all samples. However, 52.8% of CTSSs are only supported by 1-2 uniquely mapped tags (Figure S2). These CTSSs could be due to technical artifacts or the stochastic transcription, which is the main source of significant cell-to-cell variations at mRNA levels (Raj and van Oudenaarden 2008). To minimize the false CTSSs, we only considered those with TPM (tags per million) >= 0.1 for further analysis (on average, supported by at least three uniquely mapped tags). The number of qualified CTSSs ranges from 315,546 to 511,937 across the nine growth conditions, with a median of 395,182. Combination of CTSSs obtained from nine growth conditions yielded 925,804 unique CTSSs, which doubles the CTSSs identified by a single growth condition (Figure 1A and Table S2), supporting that it is necessary to examine more growth conditions to obtain a more complete TSS maps in yeast. Even though we used a conservative threshold, the number of TSSs identified in yeast from this study is significantly more than any previous study. For instance, the TIF-Seq analysis, which generated 1.88 million tags from two growth conditions, identified 227,021 TSSs supported by at least two sequencing reads (Pelechano et al. 2013). TSS identification using modified 5’RACE obtained 225,563 CTSSs with TPM >= 0.1 (Malabat et al. 2015). In the fission yeast *Schizosaccharomyces pombe*, which has a similar genome size (12.61Mb) as *S. cerevisiae*, CAGE analysis only identified 93,736 CTSSs supported by single CAGE tag (Li et al. 2015).

**Table 1.** A list of growth conditions examined by this study

| Growth Condition | Abbreviation | Description |
| --- | --- | --- |
| Cell division arrest-α factor | Arrest | 2.5mM for 45 minutes; add another 50 μL to 25mL yeast for another 30 minutes |
| Log phase | YPD | YPD medium (2% peptone, 1% yeast extract, 2% glucose) to log phase |
| DNA damage | DD | 1 mM MMS for 1 hr |
| Diauxic shift-After | DSA | YPD medium for 48h |
| Carbon source-Galactose | Gal | YP medium with 2% Galactose |
| Fermentation-16% Glucose | Glc | YP medium with 16% Glucose |
| Oxidative stress-H<sub>2</sub>O<sub>2</sub> | H<sub>2</sub>O<sub>2</sub> | Add H<sub>2</sub>O<sub>2</sub> to early-log phase cells for a final concentration of 0.20 mM, 30 mins |
| Heat shock | HS | Heat Shock from 30°C to 37°C, 1h |
| Osmotic stress-NaCl | NaCl | Add NaCl for a final concentration of 1M for 45 minutes |

**Figure 1.**
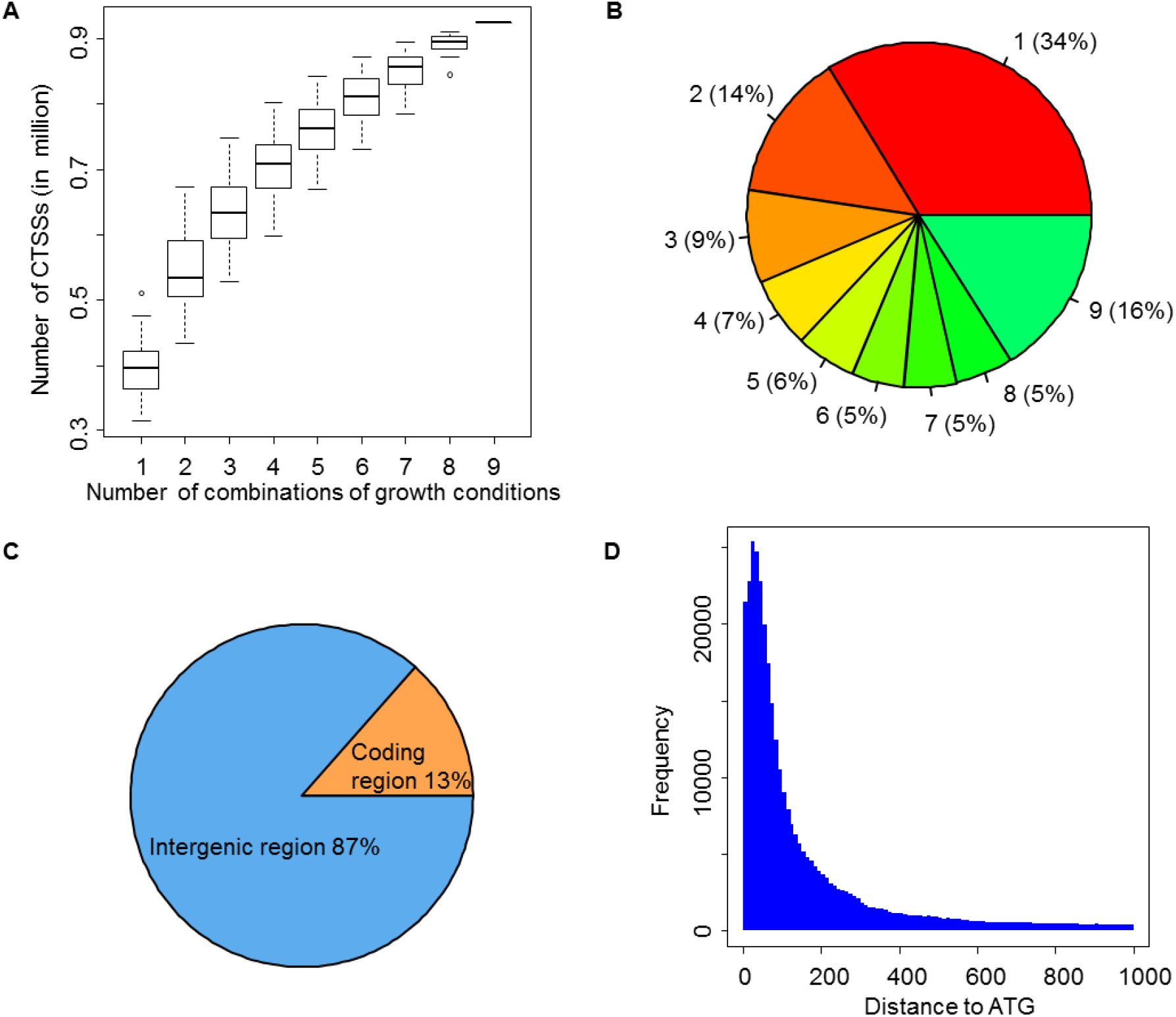
Pervasive and dynamic transcription initiations in *S. cerevisiae*. (A) The number of CTSS identified by different combinations of growth conditions. The x axis indicates all possible combinations of x number of growth conditions. (B) The distribution of CTSSs based on the numbers of growth conditions in which their activities are detected. Only 16% of CTSSs has detectable activities across all nine examined conditions. Numbers in each part of the pie chart represent the number of growth conditions. (C) Most of CAGE tags (87%) were mapped to the intergenic region. (D) A histogram shows the distribution of distance between CTSSs and translation start codon (ATG) of ORFs. CAGE tags are highly enriched in the region of 30-40 nt upstream of ATG.

As expected, most CAGE tags (87%) were mapped to the intergenic regions (Figure 1C), supporting that most transcription is initiated from non-coding regions. The CTSSs are highly enriched within 200 bp upstream of translation start codon of ORFs. The distribution of CAGE tags forms a sharp peak at ~30-40 nucleotides (nt) upstream of the ORF (Figure 1D). Therefore, the most common size of the 5’untranslated region (5’UTR) of mRNA transcripts in yeast is around 30 nt, which is probably the size optimal for binding of 40S ribosomes. It worth noting that only a small portion of CTSSs (16%) have detectable activity in all nine examined conditions. Transcriptional activity of most CTSSs was detected in part of growth conditions, and 34% of them are active in only one growth condition (Figure 1B), suggesting a highly dynamic activity of TSS in response to environmental cues.

### Identification of core promoters and improvement of genome annotation

The core promoter was typically defined as a stretch of contiguous DNA sequence encompasses the TSSs (Butler and Kadonaga 2002). The availability of multiple quantitative TSS maps allowed us to infer a more complete and accurate maps of core promoters in yeast. We used a hierarchical approach to infer core promoters by integrative analysis of the TSS maps obtained from the nine growth conditions (see Methods and materials). In brief, we first identify tag cluster (TC) from each TSS map by clustering neighboring CTSSs separated by < 20 nt. The cumulative distribution of each TC was calculated to determine the positions of the 10^th^ and 90^th^ percentile, which was defined as the left and right boundaries of a TC. For TCs are overlapped or separated by < 50 nt across different TSS maps, they were considered as the same core promoter, and were aggregated into a single consensus cluster. A total number of 43,325 consensus clusters were inferred from the nine TSS maps, representing the largest number of putative core promoters identified in yeast so far. We then assigned the consensus clusters to pol-II transcribed genes as their core promoters based on the distance between their dominant TSS (the CTSS with highest TPM) and the boundaries of downstream genes (see Methods and materials, and Figure S3).

We noticed that many consensus clusters locate within the coding or intragenic regions. Considering the pervasive nature of transcription in eukaryotes, they could be cryptic promoters within gene bodies that provoke spurious intragenic transcription (Kaplan et al. 2003). Another possibility is that the annotation of translation start codons is inaccurate (Cliften et al. 2003; Kellis et al. 2003) or the presence of alternative translation start codons (Bazykin and Kochetov 2011). Thus, we manually examined the intragenic consensus clusters to identify potentially incorrect translation start codons or alternative translation start codons (Figure 2A-C). Because translation is generally initiated at the first AUG they encounter by ribosomes during scanning of mRNA in the 5’-to-3’ direction (Kozak 2005), we searched for the first ATG codon downstream of the intragenic consensus clusters, which is likely the correct or alternative translation start codon of the ORF. Based on the location of intragenic consensus clusters, absence/presence of upstream intergenic consensus clusters, and the relative transcriptional activity between intergenic and intragenic clusters, we classified ORFs into three categories (I, II, III), representing different types of gene annotation revisions (see Methods and materials). In summary, we suggested new translation start codons for 50 genes Category I ORFs, 80 Category II ORFs, and suggested alternative translation start codons for Category III 47 ORFs (Figure S3, and Table S3), improving the annotation of the coding region of genes for *S. cerevisiae*.

**Figure 2.**
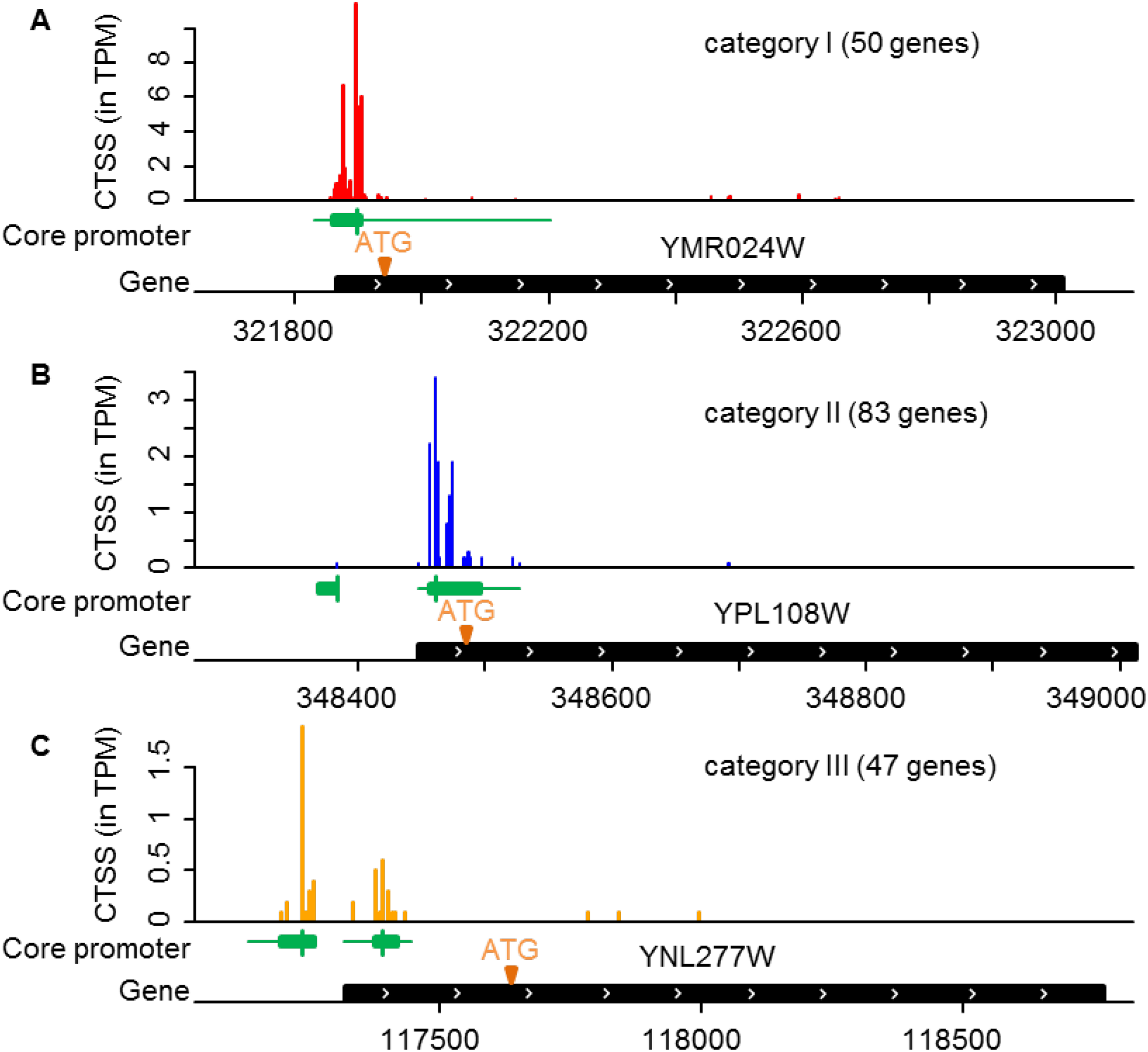
TSS maps improve yeast genome annotation. (A-B) The translation start codons of 130 genes were corrected based on location of core promoters, including both categories I (without intergenic consensus clusters) and II (with relatively low activity intergenic consensus clusters). (C) Alternative start codons were suggested for 47 category III genes. In each example, the first track is quantity of CTSS aligned to reference according to genomic coordination. The second track (green shape) represents the predicted core promoters which were generated from clustering CTSSs. Green box indicates the region enriched with the number of CTSS from 10^th^ to 90^th^ quantile. Vertical line represents the dominant CTSS in each core promoter. The third track displays the annotated gene.

Based on the revised ORF boundaries, we have assigned 11,462 consensus clusters for 5,954 ORFs as their core promoters, including 5,554 verified ORFs and 400 dubious ORFs (Table S4). Therefore, our study inferred core promoters for 95.8% of 5,797 verified protein-coding genes (assembly R64-21-1). These ORF-associated consensus clusters contain 88.5% of all uniquely mapped tags. We also assigned 255 consensus clusters, accounting for 4.9% of uniquely mapped tags, to 92 non-coding genes, such as snRNAs and snoRNAs (Table S4). Furthermore, a total number of 555, 1,944, and 370 consensus clusters were assigned to the predicted CUTs, XUTs, and SUTs respectively (Table S4) (Xu et al. 2009; van Dijk et al. 2011). The remaining 28,741 consensus clusters are not associated with any pol-II transcribed genes based on our criteria (Table S4). Most of these unassigned clusters have low transcriptional activity, and only include 5.7% of uniquely mapped tags. The presence of a large number of low-activity consensus clusters might reflect the prevalent stochastic or cryptic transcription initiation in the yeast genome. However, the possibility that they are the core promoters of unknown pol-II transcribed genes cannot be excluded, so our consensus cluster data are valuable for future studies to identify novel genes or transcripts in yeast.

### Dynamic activity of core promoters in response to environmental cues

Our comparative analysis of TSS maps unraveled a highly dynamic transcription initiation landscape in the yeast genome in response to changing environments. To better characterize the functional significance of dynamic transcription initiation, we divided the yeast protein-coding genes into two groups based on the number of core promoters assigned to them. If a gene contains only one core promoters, it was classified as single-core-promoter gene, and 44% of genes belong to this group (Figure 3A, Table S3). The remaining 56% of genes have two or more core promoters, which were classified as multi-core-promoter genes. Interestingly, the proportion of multi-core-promoter genes in yeast is similar to human, in which 58% of genes have at least two core promoters (Carninci et al. 2006). Unlike the human genome, the *S. cerevisiae* genome is highly compact, and only 30% of the genome are intergenic regions. The average length of the intergenic sequence in yeast is much shorter than that of the human genome (2.2 kb vs. 71 kb). Therefore, despite the huge discrepancy in the size of intergenic sequences, the prevalence of multi-core-promoter genes is similar between compact and large genomes.

**Figure 3.**
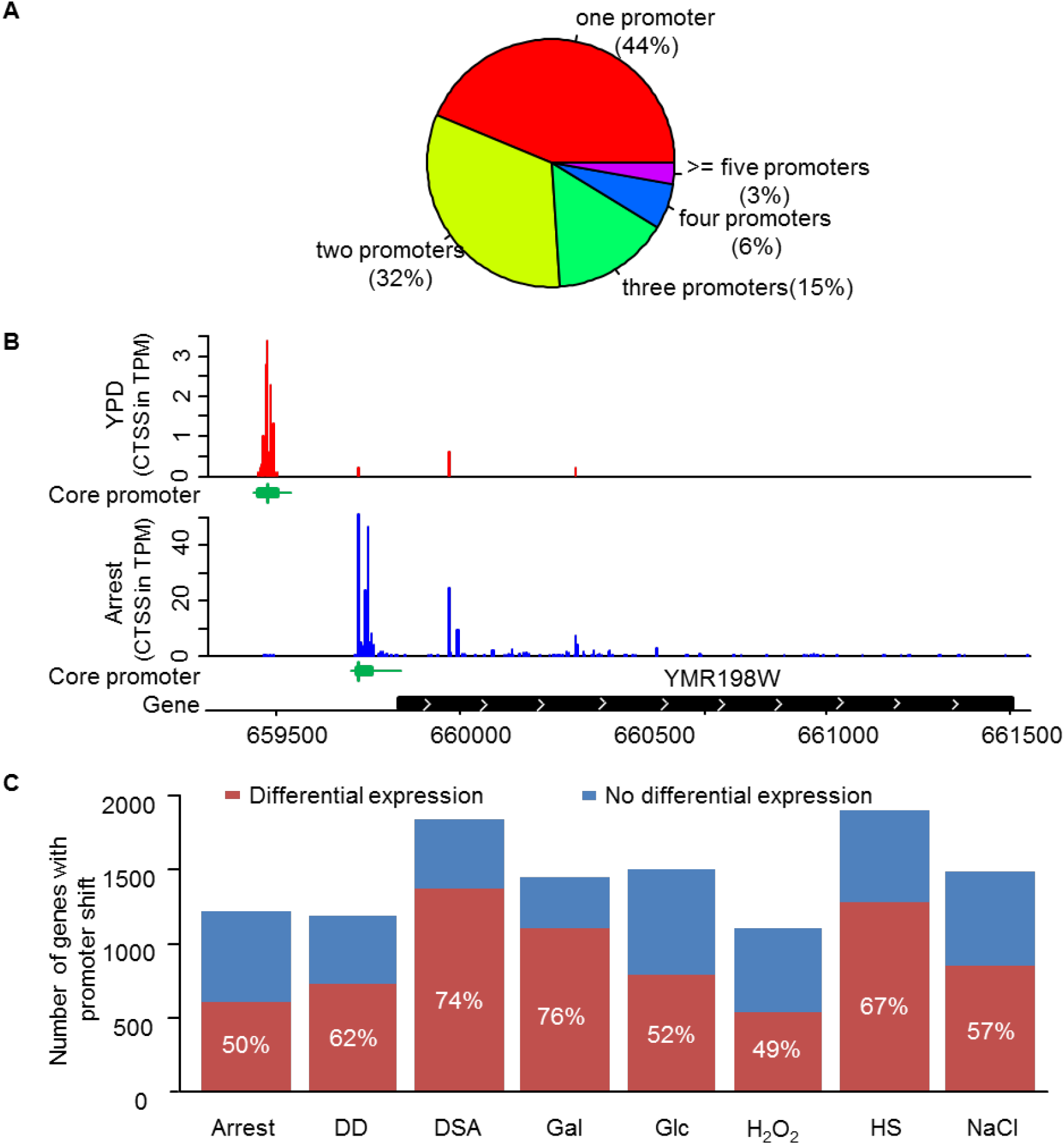
Promoter shift associates with differential gene expression. (A) Most of genes contain more than one core promoter. (B) An example of promoter shift under two environmental conditions. (C) Most of promoter shifts are associated with differential expressed genes in response to environmental cues. Labeled numbers in the bar chart show the percentage of genes encountered promoter shift.

Our comparative analysis revealed that in most multi-core-promoter gene, the proportion of CAGE signals between core promoters are significantly different in response to environmental cues, suggesting the prevalence of alternative usage of core promoter, or core promoter shift (see Methods and Materials, Table S5). Noticeably, it is also widespread that core promoter shift is coupled with gene differential expression. For instance, the *CIK1* gene (YMR198W), which encodes kinesin-associated proteins involving in controlling both the mitotic spindle and nuclear fusion during mating (Page and Snyder 1992; Shanks et al. 2001), has two core promoters located ~300 nt from each other. Transcription of *CIK1* in yeast cells grown in rich medium (YPD) is exclusively initiated from the distal core promoter and no transcription activity was detected from the proximal core promoter (Figure 3B and Table S5). Upon exposure to the mating pheromone α factor (cell arrest), almost all transcription initiation of *CIK1* switched to the proximal one (Figure 3B). Meanwhile, the transcription level of *CIK1* is significantly increased (from 12.8 TPM to 200.3 TPM). The upregulation of *CIK1* in response to *α* factor treatment is consistent with a previous study based on Northern blot (Kurihara et al. 1996). This switch of core promoter by *CIK1* is also supported by a low-throughput study using 5′RACE and the shift produced a shorter Cik1p isoform that lacks 34 AA at N-terminus (Benanti et al. 2009). Because the N-terminus of Cik1p is important for its nuclear localization and it contains a sequence that is necessary for ubiquitination, the shorter Cik1p isoform is stable. Therefore, this case suggests that core promoter shift may have a significant impact on gene expression and protein function.

To measure the degree of prevalence of core promoter shift across different growth conditions, we quantified a gene’s core promoter shift based on the changes of CAGE tag distribution between core promoters for all multi-core promoter genes (see Methods and Materials). Using the core promoter activity in yeast cells grown in YPD as a control, we found that 2,833 of 3,349 (85.6%) multiple-core-promoter genes have experienced a significant shift of core promoter activity in at least one treatment, supporting that core promoter shift in response to changing environments is widespread in yeast. Depending on growth conditions, 48.9% to 76.1% of genes with core promoter shift are coupled with significant gene differential expression (Figure 3C). Therefore, similar to the regulated alternative core promoter usage in different tissues in human and mouse, condition-specific transcripts are also commonly generated by alternative usage of core promoters in the unicellular eukaryotic organisms. In short, our results revealed that core promoter shift in yeast is widespread, and most shifts are associated with gene differential expression, suggesting that shift of core promoters might function as an important mechanism for fine-tuning of gene expression in yeast.

### Two classes of core promoters in *S. cerevisiae*

The regulated transcription initiation was mostly characterized at the gene level in previous studies. Considering that most genes contain multiple core promoters and the alternative usage of core promoters are prevalent, it is more informative to characterize the dynamic of transcription initiation at the core promoter level. Among the 11,462 core promoters assigned to protein-coding genes, only 55% of them (6,251) have detectable transcriptional activities under all nine examined conditions (Figure 4A). The transcriptional activity of 17% core promoters can only be detected under one growth condition, suggesting a strong condition-specificity (Figure 4A). Based on their transcription activities across nine growth conditions, we classified yeast core promoters into two classes: constitutive core promoter and inducible core promoter (Table S4). If transcription initiation constitutively occurs from a core promoter in all examined growth condition, it is defined as constitutive core promoter. Only 5,211 (45%) core promoters belong to the constitutive class. If the transcriptional activity of a core promoter can only be detected under one or some of the examined growth conditions, it is classified as inducible core promoters, which include 55% of core promoters (Figure 4A).

The distributions of the two classes of core promoters are significantly different between the single-core-promoter genes and multi-core-promoter genes (Figure 4B-C). Specifically, 88% of core promoters in single-core promoter genes are constitutive core promoters, while only 45% of core promoters in multi-core-promoter genes are constitutive core promoters. This observation suggests that most single-core-promoter genes tend to be constitutively expressed in yeast regardless of growth environments. In contrast, in the multi-core-promoter genes, most of their core promoters are only active under specific conditions. In eukaryotes, constitutively expressed genes are usually required for the maintenance of basic cellular function, which were also called housekeeping genes (Zhu et al. 2008). Consistently, based on our Gene Ontology (GO) enrichment analysis (Figure S4), single-core promoter genes are generally enriched in these housekeeping genes, which are related to cellular component and organelle organization, macromolecule, cellular, and protein localization. In contrast, multi-core promoter genes are more enriched in gene regulation process, such as regulation of biological process, regulation of cellular process, biological regulation, etc. (Figure S4).

**Figure 4.**
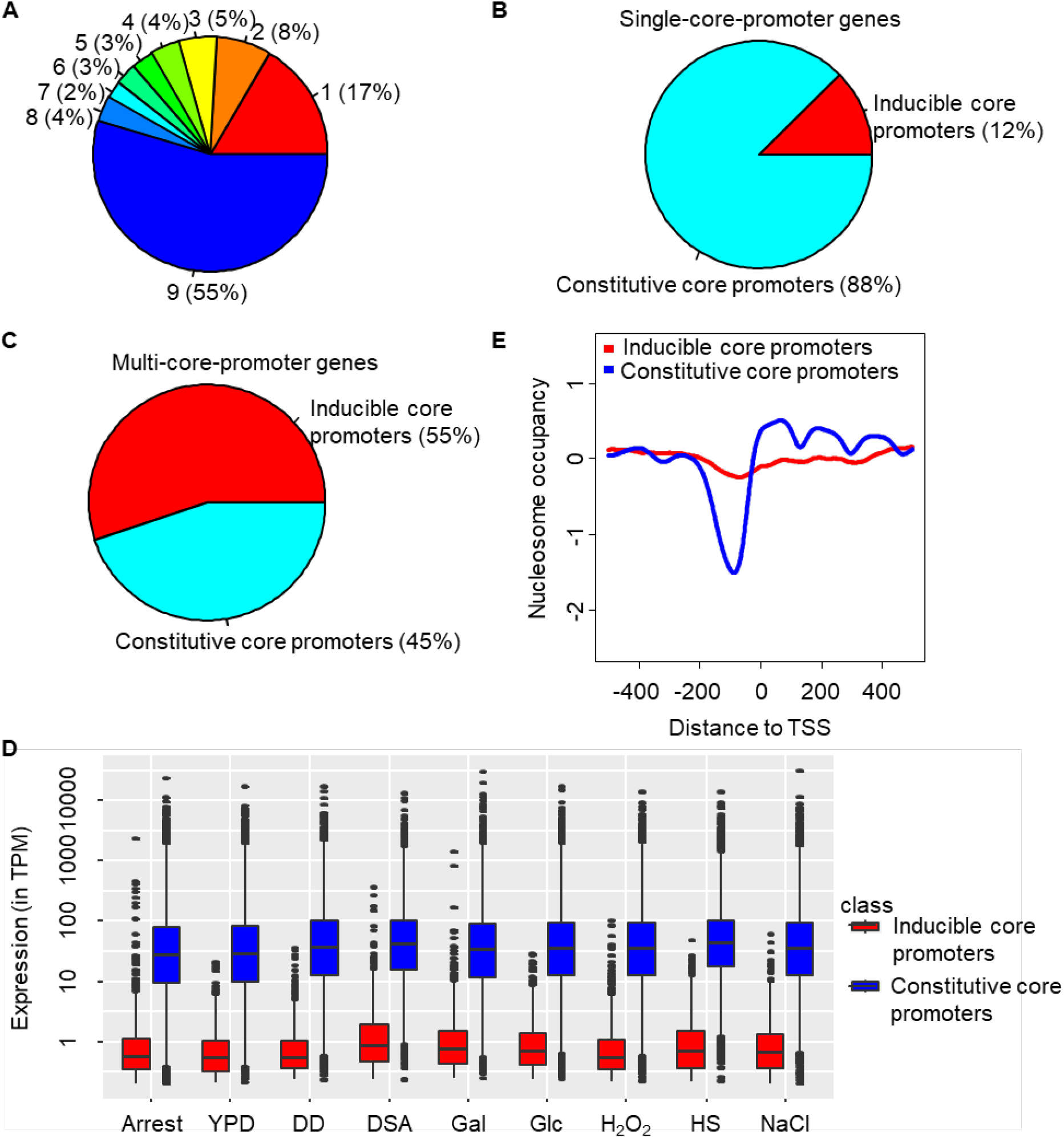
Distinct structures and expression patterns between constitutive and inducible core promoters in *S. cerevisiae*. (A) Pie chart shows the activity of core promoters under different numbers of growth conditions. Numbers in the pie chart represent the number of combinations of growth conditions. Over half of core promoters (55%) which were expressed under all nine conditions are constitutive core promoters, and the others are inducible core promoters. (B) Majority of core promoters (88%) in single-core-promoter genes are constitutive core promoters. (C) Most core promoters (55%) in multi-core-promoter genes are inducible core promoters. (D) Constitutive core promoters are associated with much higher transcriptional activity than inducible core promoters. (E) In rich median (YPD), constitutive core promoters are associated with depleted nucleosome occupancy while inducible core promoters tend to be occupied by nucleosome.

Another distinct pattern between the two types of core promoters is their transcriptional abundance. At a genome-scale, the transcriptional activity from constitutive core promoters is significantly stronger than that of inducible core promoters in all nine growth conditions (Figure 4D). We speculated that the different transcriptional abundance between the two types of core promoters is due to different nucleosome positioning patterns, which were shown to have major impacts on transcriptional activity (Jiang and Pugh 2009; Nocetti and Whitehouse 2016). The eukaryotic DNA is coiled around a core of histones which forms nucleosomes. If nucleosome is present in the core promoter region, it becomes an obstacle for transcription initiation. Thus, the activation of transcription from these core promoters requires alteration of chromatin structure by ATP-dependent nucleosome sliding (Shen et al. 2000; True et al. 2016) or histone modification (Shilatifard 2006). To determine whether inducible core promoters are generally occupied by nucleosomes to allow condition-specific activation, we compared the chromatin structure near the core promoters (± 500 nt of dominant TSS) between the two classes of core promoters using the nucleosome occupancy data obtained from (Field et al. 2009). We observed a nucleosome-free region (NRF) immediately upstream of TSS in constitutive core promoters (Figure 4E, Figure S5-6). In contrast, the inducible core promoters generally are occupied with nucleosomes in the same region. Most of inducible core promoters are inactive in yeast cells grown in YPD. We also observed a more depleted nucleosome occupancy upstream of TSS in the inducible core promoters that are active under YPD growth condition than those are inactive (Fig. S5), supporting that nucleosome occupancy pattern plays a determining role in controlling the transcriptional activity of a core promoter. Therefore, because of the difference in chromatin structure, different mechanisms of transcription activation are likely involved in the two types of core promoters. It was proposed that nucleosome positioning is largely determined by the intrinsic property of nearby DNA sequences (Kaplan et al. 2009; Tirosh et al. 2010). It is possible that the different genomic context might underlie the distinct chromatin structure between the two types of core promoters.

### Distinct promoter shape between inducible and constitutive core promoters

Transcription can be initiated at precise positions or a disperse region, which form a continuum of shape of core promoters from sharp to broad shape (Carninci et al. 2006; Hoskins et al. 2011). The promoter shape is generally conserved between different species (Carninci et al. 2006; Main et al. 2013), and different signatures of evolution have been observed between broad and sharp core promoters (Schor et al. 2017), supporting an important but distinct functional role between promoters with different shapes. Similar to metazoan species, the spatial distribution of CAGE signals varies substantially among core promoters in *S. cerevisiae*, spanning a range of shapes from peaked to broad (Figure 5A). To characterize the shape of yeast core promoters and to determine the extent to which inducible core promoters differ from constitutive core promoters in promoter shape, we developed a new metric to describe promoter shape, called Promoter Shape Score (PSS, see Methods and Materials). PSS integrates the observed probability of tags at each TSS within a core promoter and the quantile width of a core promoter. The main improvement of PSS over previous promoter shape estimation method Shape Index (SI) (Hoskins et al. 2011) is that SI does not take into consideration the distances between TSSs, which determine the promoter width. Without integrating promoter width factor, SI does not distinguish the difference between two promoters if tags are discontinuously distributed.

**Figure 5.**
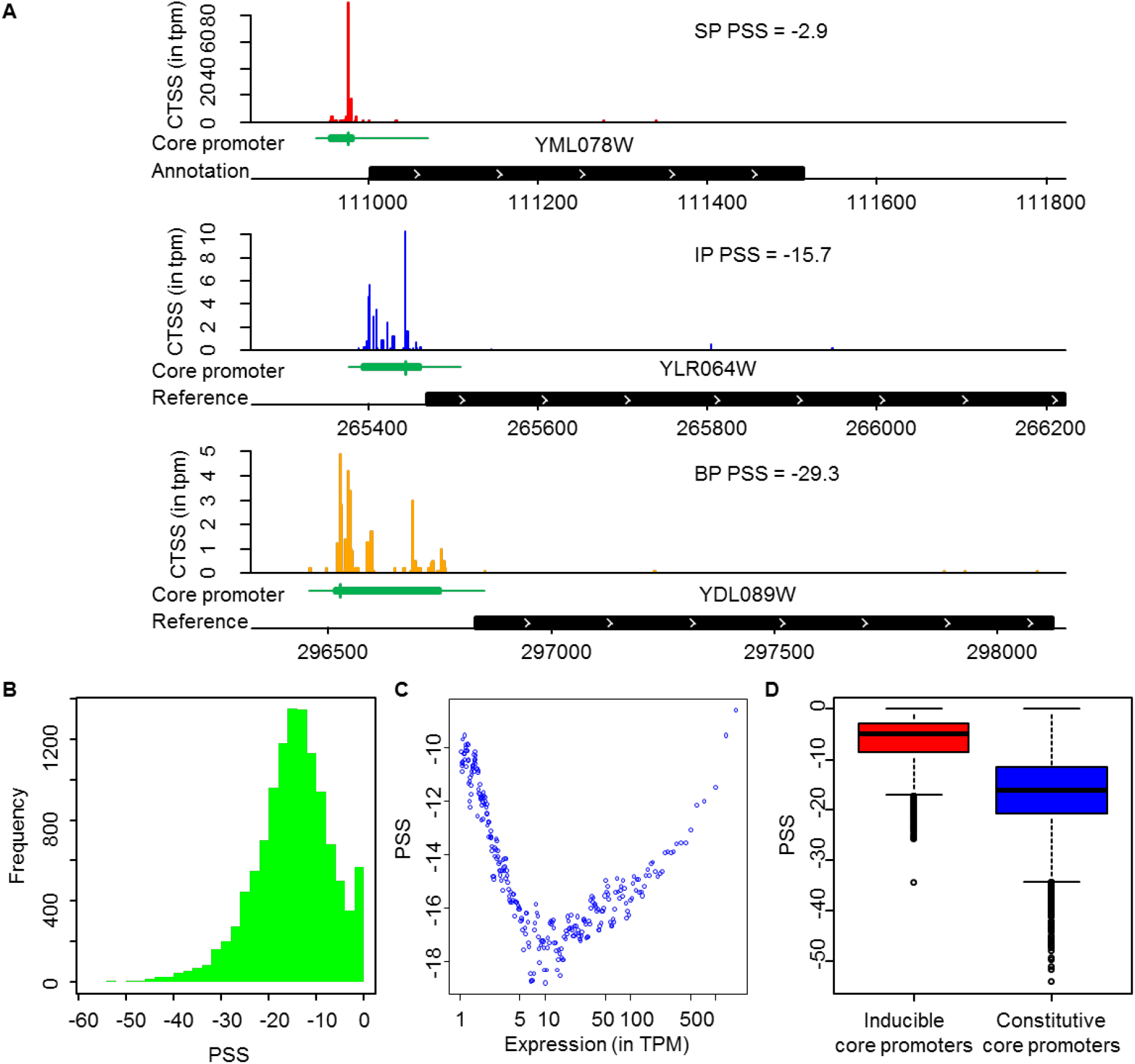
Core promoter classification. (A) Examples of sharp, intermediate and broad core promoter in *S. cerevisiae*. Core promoters with PSS greater than -10 were classified as sharp core promoters (SP), smaller than -20 as broad core promoters (BP), and the others as intermediate core promoters (IP). (B) Histogram shows the continuous distribution of PSS values in *S. cerevisiae*. (C) Lowly and highly expressed core promoters tend to be sharp (higher PSS score). Dot plot was drawn based on a sliding window analysis after sorting all core promoters by transcription abundance (TPM). Window size is 200 core promoters and moving step is 40 core promoters. Each dot presents the median values of PSS of 200 core promoters. (D) Box plot of PSS values of inducible core promoters and constitutive core promoters.

Based on our algorithm of PSS, the sharpest core promoter has a PSS value of 0, which means that all transcription initiation of a core promoter occurs from a single TSS (also called singleton). The PSS value decreases when the number of TSSs increases and/or more even transcription from different TSSs. The PSS values of all core promoters of protein-coding genes largely follow a Gaussian distribution (-14.8±8.07). We noticed that there are more core promoters with a PSS close to 0 than expected (Figure 5B). This is due to the presence of many singleton core promoters. As shown in Fig. 5B, PSS values form a peak in the range of from -20 to -10. Therefore, based on PSS values, we classified core promoters into three groups: sharp core promoter (SP) with PSS > -10; broad core promoters (BP) with PSS <= -20; and the rest are considered as intermediate shape core promoters (IP) (Figure 5A).

To determine the relationships between core promoter shape and gene expression patterns in yeast, we conducted a sliding-window analysis between PSS and their transcription abundance. By plotting the median PSS and TPM values of each window of 200 core promoters, we observed a “V” shape between the two metrics (Figure 5C). In general, for core promoters with transcription abundance < 10 TPM, the core promoters become broader with increase of transcription abundance. In core promoters with the lowest activity, transcription is usually initiated from a single TSS, which form an ultra-sharp promoter (PSS = 0). The increase of transcription activity for these low-activity core promoters appears to be mainly achieved by expanding TSSs, resulting a broader promoter shape. However, for core promoters with transcription abundance > 10 TPM, the core promoters become sharper with increase of transcription abundance. It means that the increase of transcription abundance is mainly achieved by increased transcription from one or a few TSSs within these core promoter, rather than expanding the TSSs, which forms a positive correlation between transcription activity and PSS.

The PSS values of constitutive core promoters are significantly lower than that of inducible core promoters (p < 2.2×10^-16^, Student’s t-test, Figure 5D). Of 5,211 inducible core promoters, 4,267 of them were classified as sharp promoters, and only 42 are in the broad class. In contrast, among the 6,251 constitutive core promoters, 1,111 are sharp promoters and 1,732 are broad promoters. This is because most of inducible core promoters have TPM < 1, and constitutive core promoters have a broader range of transcriptional abundance (Figure 4E). As shown in Figure 5C, core promoters with TPM < 1 have high PSS values, or sharp shape. A previous study in *Drosophila* showed that sharp core promoters are more likely to have restricted tissue-specific expression, while broad core promoters to have a constitutive temporal expression pattern in *Drosophila* (Hoskins et al. 2011). Similar to *Drosophila*, inducible core promoters have restricted condition-specific expression and have a sharper shape than constitutive core promoters, support that the different regulatory mechanism of transcription initiation of the two classes of core promoters are conserved between yeast and animals.

### Strong preference of pyrimidine-purine dinucleotides at yeast TSSs

In animals, transcription preferentially starts with pyrimidine-purine (PyPu) dinucleotide, that is a purine at position +1 (TSS), and a pyrimidine at position –1 (Burke and Kadonaga 1997). Here, we examined the TSS dinucleotide preference in *S. cerevisiae* and found that 86.7% of CAGE tags were mapped to PyPu dinucleotide at position –1,+1, supporting strong evolutionary conservation of PyPu dinucleotide preference in eukaryotes. We then investigated whether inducible and constitutive core promoters have different dinucleotide preference. Using the TSS with the highest transcription activity (dominant TSS) as the representative TSS for each core promoter, we obtained the consensus sequence around TSS (±10 nt) for each type of core promoters. As shown in Fig. 6A-C, PyPu dinucleotide at position –1,+1 of TSS are highly enriched in both types of core promoters. The most preferred PyPu dinucleotide is CA, following by TA and TG (Figure 6C). However, constitutive core promoters have a stronger preference for PyPu dinucleotide at TSS than inducible core promoters (95.0% vs. 76.8%, Figure 6C). Furthermore, the constitutive core promoters have a much stronger preference of CA dinucleotides than the inducible core promoters. In addition to the PyPu dinucleotides, the DNA sequences surrounding the dominant TSSs in yeast is enriched of A, especially at position -8 in both types of core promoters, which is a pattern that is not present in other species (Figure 6A-B). However, the frequency of A at position -8 in constitutive core promoters is higher than inducible core promoters (72.6% vs. 58.8%), suggesting a different sequence preference of transcription initiation between the two types of core promoters.

**Figure 6.**
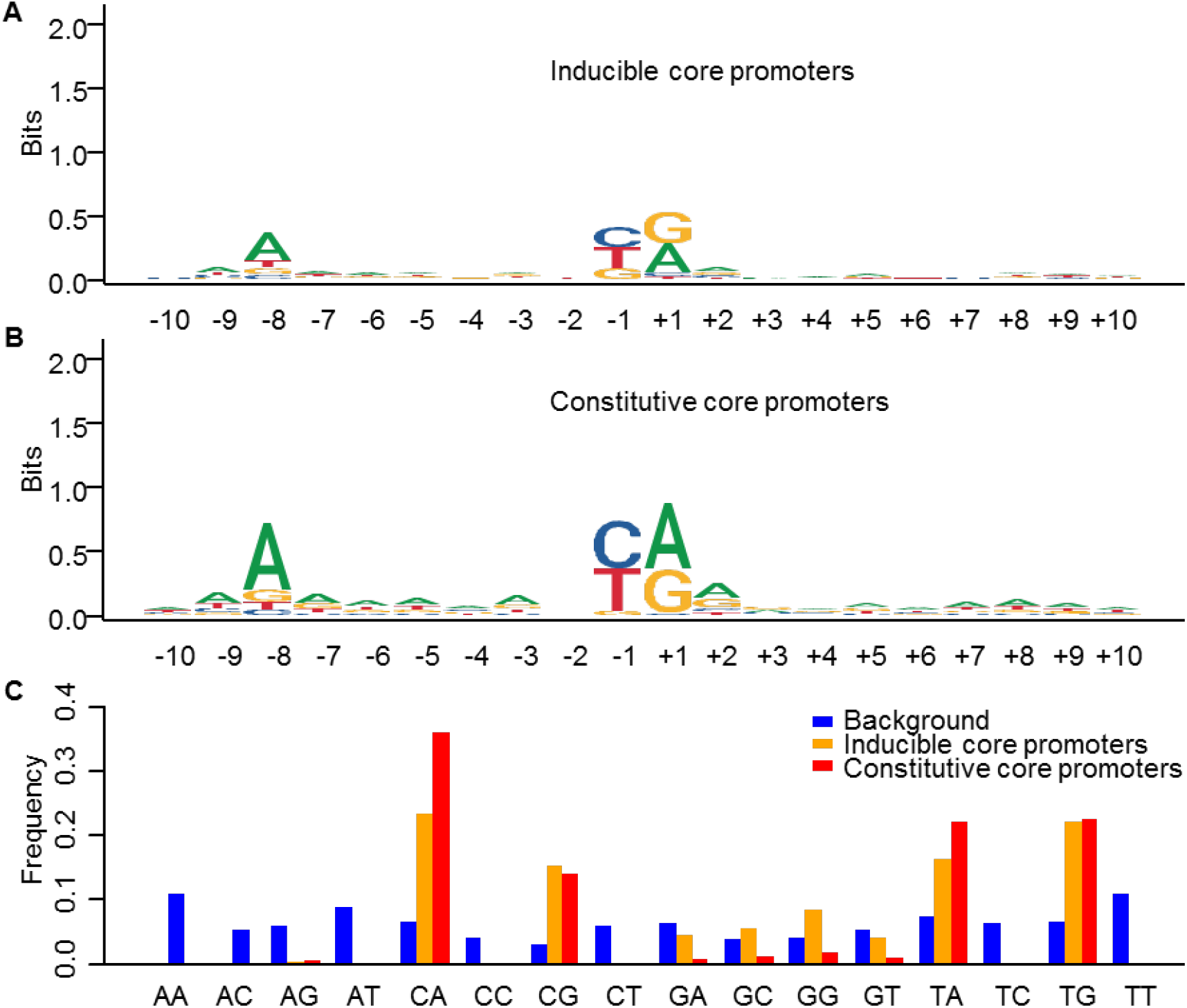
Initiator motif and dinucleotide preference of core promoters in *S. cerevisiae*. Sequence logo of the consensus sequence of 20 nt surrounding the dominant TSS of (A) inducible core promoters and (B) constitutive core promoters, which likely represents the Initiator element in yeast. (C) TSSs in *S. cerevisiae* have a strong preference of pyrimidine-purine dinucleotide at [+1,-1] positions. CA is the most frequent TSS dinucleotide, followed by TA and TG.

### Different DNA motifs between inducible and constitutive core promoter

We sought to identify overrepresented DNA motifs in or near the inducible and constitutive core promoters to explore their different references of sequence context. We performed *de novo* motif discovery for the surrounding sequences of these core promoters (from -100 to +50 of TSS). Predicted DNA motifs from each class of core promoters were compared with known binding motifs in *S. cerevisiae* to identify possible matches (Zhu and Zhang 1999; MacIsaac et al. 2006). Among the top enriched motifs, only two are shared by the two classes of core promoters (Figure 7). One of them has a consensus sequence of “3’-TATAAA(A)AAA-5”, has significant similarity with the cononical binding sites of TATA binding protein (TBP), the TATA-box (Table S6). TBP is a subunit of the TFIID complex in eukaryotes that recruits the transcriptional machinery to the promoter. A total number of 6,843 TATA-box motifs are found near 1,269 inducible core promoters and 1,500 constitutive core promoters (-100 and + 50 nt of dominant TSS, Table S6). The percentage of TATA-box containing promoters with is virtually the same between inducible core promoter sequences (24.35%) and constitutive core promoters (24%). A total number of 2,284 protein-coding genes were found to be associated with at least one TATA-box motif, which is about twice as many as previous identified based on ChIP-chip assays of TBP (Basehoar et al. 2004). In metazoans, transcription typically initiates 25-30 bp downstream the TATA-box. However, it was believed that transcription in *S. cerevisiae* initiate from a wide range of 40–120 bp downstream of the TATA box, lacking a strictly defined distance (Smale and Kadonaga 2003). In this study, with more accurate TSS maps generated by this study, we found that the distribution of TATA-box forms a sharp peak around 65 bp upstream of TSS in both inducible and constitutive core promoters (Figure 8). Therefore, our data suggest that the location of TATA-box in *S. cerevisiae* is also strongly confined, similar to that of metazoans. The main difference is that the preferred distance of between TATA-box and TSS in yeast is about 40 bp longer than that of metazoans, suggesting a different mechanism of transcription initiation in yeast. It has been proposed that in yeast PIC assembles at a TATA box, and pol-II searches the template strand for acceptable TSSs, which is called the scanning model (Giardina and Lis 1993). Here, we showed that there is still a strong preference of distance between TATA-box and TSSs in yeast, suggesting that the distance is probably still an important factor when PIC searches for acceptable downstream TSSs.

**Figure 7.**
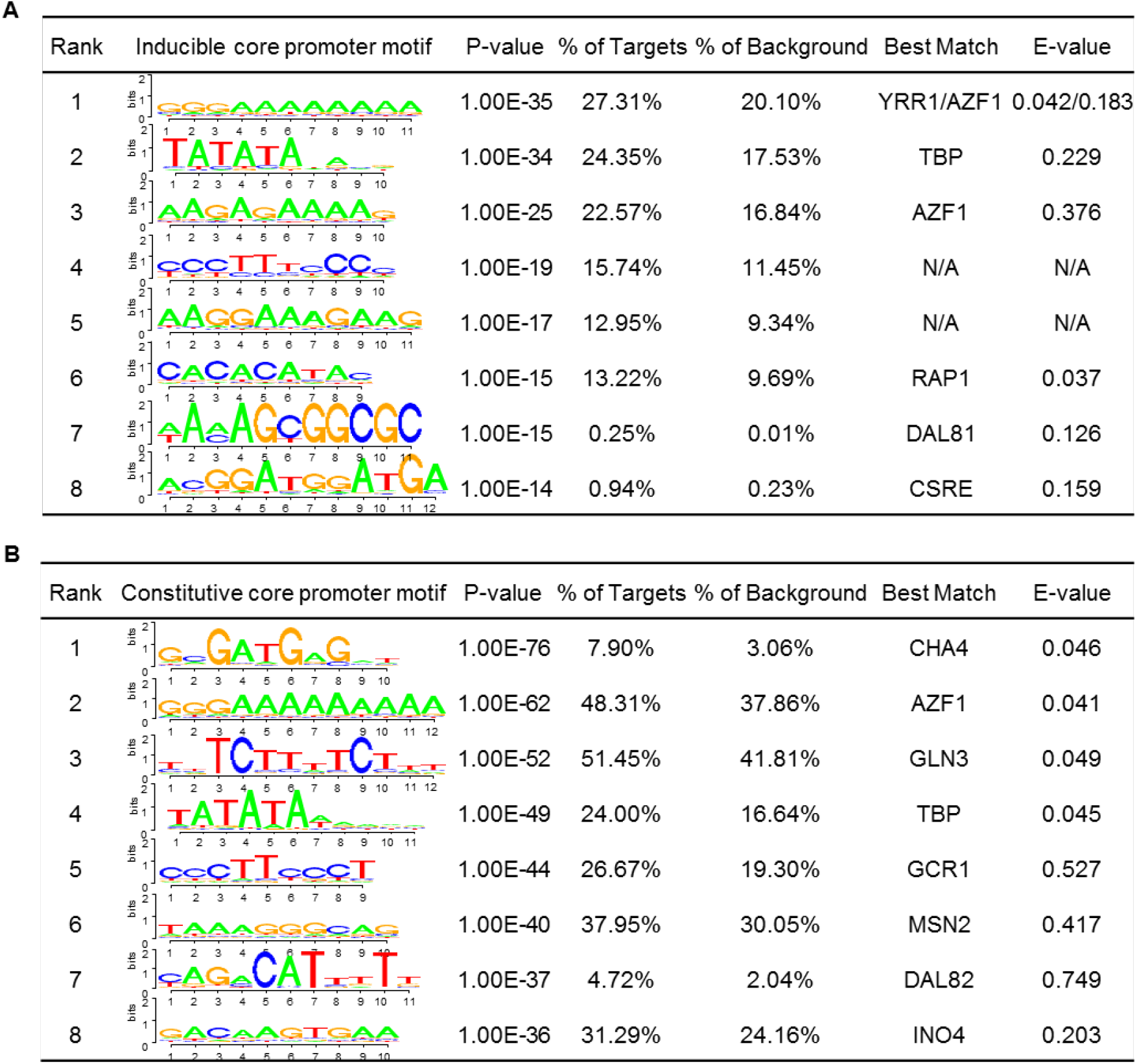
Predicted core promoter motifs in *S. cerevisiae*. (A). The top enriched motifs present in the inducible core promoter sequences. (B) The top enriched motifs present in the constitutive core promoter sequences. These promoter motifs were predicted by *de novo* discovery approach for the 150 bp sequence surround the dominant TSS (-100 and +50 bp) in each core promoters.

**Figure 8.**
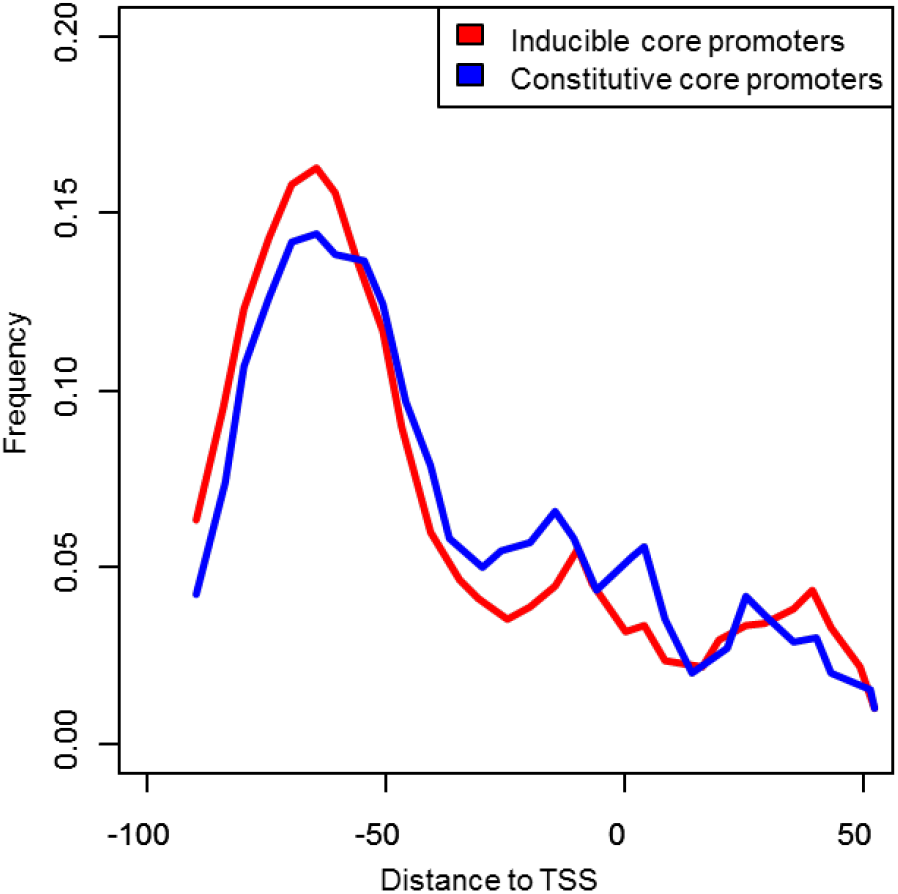
TATA box is highly enriched around 65 bp upstream of TSS in both inducible and constitutive core promoters in *S. cerevisiae*. The distribution of TATA box was inferred by sliding window analysis with a window size of 10 bp and moving step of 5 bp.

The other shared motif, with a consensus sequence of GGGAAAAAAAA, is present in nearly 50% of core promoters. It is most similar to the binding motif of YRR1 or AZF1, both of which are zinc-finger transcription factors. YRR1 is involved in multidrug resistance (Cui et al. 1998), while AZF1 is involved in diauxic shift and response to hypoxia (Newcomb et al. 2002). As these two transcription factors are only in specific cellular processes, this motif is unlikely the binding sites of YRR1 or AZF1, despite the high similarity. We also noticed that two highly enriched motifs, with consensus sequences of CCCTTTCCCC and AAGGAAAGAAG (Fig. 7A), do not have a significant match with any known motif sequence. As only a small portion of eukaryotic genes have the TATA-box, the known binding sites for TBP. It remains obscure about what binding motifs are bound by general transcription factors in the TATA-less genes. We speculated that some of these high-frequency motifs, might be the binding sites of general transcription factors. Future studies can focus on these motifs to identify their binding proteins and their roles in gene transcription.

## DISCUSSIONS

In this study, we generated a comprehensive quantitative TSS maps for the important eukaryotic model organism *S. cerevisiae* at an unprecedented resolution and depth. Our study unravels the highly pervasive and dynamic nature of transcription initiation in yeast. Pervasive transcription has been known in mammalian genomes (Consortium et al. 2007; Kapranov et al. 2007) as well as in *S. cerevisiae*, but there is still “blank space” to be filled by the right study (Libri 2015). We found that the transcription in yeast can be initiated from much more genomic positions than previously recognized. In the12-million bp yeast genome, there are over 4 million TSS positions supported by at least one CAGE tag, and about 1 million TSS positions were supported by multiple CAGE tags, which is still much more than the previously identified number. The biological significance of pervasive transcription is unclear and controversial (Kapranov et al. 2007). Nevertheless, our study not only supports that most eukaryotic genome is used to transcribe RNA, and it also reveals that transcription can also initiate in a large portion of yeast genome.

The increase of sequencing depth and examined growth conditions allowed us to identify the TSSs and core promoters for lowly-expressed or condition-specific expressed genes. Based on nine different TSS maps, we were able to determine the TSS and core promoters for 96% of verified ORFs in *S. cerevisiae*. In addition to determining the 5’boundary for majority of protein-coding genes, we also we revised the annotation of translation start codon for 180 ORFs. However, there are still many TSS clusters were not assigned to any known gene features. Transcriptional activities of most of these clusters are not intensive in examined conditions. The presence of large numbers of these low-activity TSS clusters is likely the consequence of the pervasive nature of transcription initiation. However, we cannot exclude the possibility that some of them could be the functional core promoters of unidentified protein-coding genes, or non-coding RNA genes. Future comparative and functional genomics studies may focus on these core promoters to infer their functional significance.

Comparative analysis of quantitative core promoter maps also allowed us to identify two types of core promoters (inducible and constitutive) and to characterize the differences in activity, stuture and sequence underlying different types of core promoters in yeast. The constitutive core promoters tend to have higher transcriptional activities than inducible core promoters in all growth conditions examined (Fig. 4D), a more nucleosome-depleted region upstream of TSS (Fig. 4C), a broader promoter shape (Fig. 5C), and stronger preferences of PyPu dinucleotides at TSSs (Figs. 6-7). These observations suggest the presence of two distinct regulatory mechanisms of transcription initiation in the unicellular organism.

One of the most interesting findings of our study is widespread core promoter shift coupled with gene differential expression in response to environmental cues in yeast. Alternative promoter usage in different cell types or tissues has been observed in mammals and fruit fly *D. melanogaster* (Davuluri et al. 2008; *Batut* et al. 2013). However, how different core promoters in multi-core promoter genes are regulated in response to environmental cues have not been systematically systematically investegated. Our data showed that most of yeast genes have at least two core promoters. The activity switch of different core promoters of a gene is prevalent across different growth conditions. Therefore, it appears that alternative core promoter usage is a conserved trait in both single- and multiple-cellular organisms. Most strikingly, we found that most of core promoter shifts in yeast are coupled with significant differential gene expression. For microorganisms, modulation of gene expression plays a central role in the adaptation to changing environmental cues (Lopez-Maury et al. 2008). The primary driver of alternative gene expression in response to changing environments is probably the switch of different condition-specific transcription factors, triggered by signal-transduction pathways through sensing extracellular signals (Lopez-Maury et al. 2008). Repositioning of nucleosomes in the core promoter regions may also play an important role in the alternative usage of core promoters. It was found that most of gene differential expression is associated with extensive nucleosome repositioning in the gene promoters (Nocetti and Whitehouse 2016).

We speculated that the shift of core promoters might serve as a secondary control for further tuning the outcome of gene expression by influencing both transcription and translation processes. These structural differences among core promoters could influence the efficiency of transcription initiation (Kostrewa et al. 2009). In addition, as a direct consequence of core promoter shift, transcripts with various length and sequence of 5’UTR are generated. Different lengths of 5’UTR may have different mRNA folding structures, which might change their thermostability. Modulation of mRNA stability is a critical step in the regulation of gene expression. In eukaryotic cells, the decay rates of individual mRNAs vary by more than two orders of magnitude (Harigaya and Parker 2016). Furthermore, the change of 5’UTR by core promoter shift could theoretically influence translation initiation efficiency, which is the rate at which ribosomes access the 5’UTR and start translating the ORF (Kudla et al. 2009; Livingstone et al. 2010). Translation initiation efficiency is highly correlated with translation efficiency (Weinberg et al. 2016), and nearly 100-fold range of translation efficiency has been observed in log-phase yeast (Ingolia et al. 2009), suggesting a significant control in yeast. Translation initiation is the main rate-limiting steps of gene expression (Pop et al. 2014). Strong secondary structure near the 5’cap might interfere with binding of the eIF4F-cap-binding complex, and structure within the 5’UTR can impede the scanning by 40S ribosome, thereby reducing the rate of protein synthesis (Ding et al. 2012). Different 5’UTR lengths may change the secondary structure near the 5’cap, which influence translation initiation probabilities (Shah et al. 2013). For instance, insertion of a stem-loop into the 5’UTR of PGK1 mRNA effectively blocks translation by preventing 40S scanning (Muhlrad et al. 1995). Our previous studies also observed some connection between 5’UTR lengths and gene expression profiles within and between yeast species (Lin et al. 2010; Lin and Li 2012), suggesting a potentially functional role of 5’UTR length in gene regulation. Therefore, we recommend that more studies will be needed to investigate the functional impacts of core promoter shift and 5’UTR length changes on gene expression, which might potentially uncover a new layer of gene regulatory mechanism.

## METHODS AND MATERIALS

### Yeast strain and growth conditions

The *S. cerevisiae* laboratory strain BY4741 (*MATa his3Δ1 leu2Δ0 met15Δ0 ura3Δ0*) was used as an experimental system to generate condition-specific CTSS maps. In the natural environments, yeast cells constantly exposed to a wide range of environmental stresses, such as temperature changes, increases in oxidative stress, imbalances in osmolarity, changes in external pH, nutrient supply, the presence of radiation and toxic chemicals. The 9 growth conditions applied in this study (Table 1) simulate these natural environmental stresses. All incubations were at 30°C, except for when heat stress when applies.

### CAGE library preparation and sequencing

Isolation of total RNA was performed using solid phase extraction (Bioline). The total RNA samples were snap-frozen in liquid nitrogen and stored at -80°C. RNA samples were quantified and evaluated for quality and using the Bioanalyzer 2100 (Agilent Technologies). 5ug total RNA sample were isolated from each sample. Two biological replicates of CAGE libraries were constructed for samples of each growth condition following the nAnT-iCAGE protocol (Murata et al. 2014) by the DNAFORM, Yokohama, Japan. In brief, RNA quality was assessed by Bioanalyzer (Agilent) to ensure that RIN (RNA integrity number) is over 7.0, and A260/280 and 260/230 ratios are over 1.7. First strand cDNAs were transcribed to the 5’ end of capped RNAs, attached to CAGE "barcode" tags. Each nAnT-iCAGE library used linkers with specific barcodes and were sequenced using Illumina NextSeq (single-end, 75-bp reads) at the DNAFORM, Yokohama, Japan. The number of reads generated from each library are listed in Table S1.

### CAGE processing, alignment, and rRNA filtering

A total number of 636 million reads were generated from the 18 samples. The sequenced CAGE tags were respectively aligned to the reference genome of *S. cerevisiae* S288c (R64-2-1) using HISAT2 (Kim et al. 2015). To avoid identification of false TSS, soft clipping option in HISAT2 is disabled by using ‘--no-softclip’. The numbers of reads mapped to the *S. cerevisiae* reference genome are provided in Table 2. The reads mapped to rRNA sequences (28S, 18S, 5.8S, and 5S) were identified from read alignments (in SAM format) using rRNAdust (http://fantom.gsc.riken.jp/5/sstar/Protocols:rRNAdust) and were subsequently removed by house-made R scripts. Tags mapped to multiple genomic regions (SAM MAPQ < 20) were excluded. Only the uniquely mapped tags were used for further analysis. All unique 5’ends of tags were identified as CAGE tag-defined TSSs (CTSSs) by house-made R scripts. The replicates of CAGE tags obtained from the same growth condition were merged. The numbers of CAGE tags supporting each CTSS were counted and normalized to tag per million (TPM) using the CAGEr package (Haberle et al. 2015) in R Bioconductor.

### Analysis of mapped CAGE tags

The CTSSs with minimum of tag per million (TPM) value 0.1 were used as input for tag clustering to infer putative core promoters. CTSSs separated by <20 bp were clustered into a larger transcriptional unit, called tag cluster (TC). Only TCs with a minimum of 0.2 TPM were used for further analysis. For each TC, we calculated a cumulative distribution of the CAGE tags to determine the positions of the 10^th^ and 90^th^ percentile, which were considered as its boundaries. TCs were first generated from each sample separately. Based on TC locations across nine samples, if two TCs are located within less than 50 bp, they likely belong to the same core promoters, so they were aggregated into a consensus cluster.

We assigned the consensus clusters to pol-II transcribed genes follow the following procedures. First, we identified the dominant CTSS for each consensus cluster. The dominant CTSS is the CTSS with largest TPM in the cluster. If a dominant CTSS is located within 1,000 nt upstream of translation start codon of the immediate downstream gene and has no overlap with the upstream gene’s coding region, this consensus cluster is assigned to the downstream gene. If the dominant CTSS of a consensus cluster is within 1,000 nt upstream of start codon of the immediate downstream gene, but it overlaps with the upstream gene’s coding region, it is assigned to the downstream gene only if it is within 500 bp. Based on these criteria, 11,462 consensus clusters were assigned to 5,954 protein-coding genes, which accounts for 90% of all predicted ORF in *S. cerevisiae*.

If a gene is not associated with any upstream intergenic consensus cluster, but has an intragenic consensus cluster located within its first half of coding regions, the downstream ATG is likely the correct start codon of this gene. Therefore, we suggested a new start codon for such genes. These genes were classified as category I and we found 50 genes belong to this group. If a gene is associated with at least one intergenic consensus cluster, we then calculated the TPM ratio between intragenic consensus cluster to the highest expressed intergenic ones (R = intragenic/intergenic). If R ≥ 2 (Figure 2B), it suggests that this intragenic consensus cluster is likely used as main core promoter for this gene, and the downstream ATG is predominantly used for this gene. Thus, for these genes, the most frequent translation start codon should be the one downstream of this intragenic Cluster. We grouped this type of genes as category II, which includes 83 genes (Table S3). If 2 > R ≥ 0.5 (Figure 2C), the intragenic core promoter might serve as an alternative core promoter of the gene. There are 47 genes that belong to this group, which was classified as category III genes (Table S3). If R < 0.5, indicating a low relative transcription activity of intragenic core promoter, they are likely the cryptic promoters.

We then assigned the remaining consensus clusters to pol-II transcribed non-coding genes, including 50 LTR retrotransposons, 77 snoRNA genes, six snRNA genes, one telomerase RNA gene. As the annotation of these non-coding genes refer to the transcribed regions, thus we only assigned consensus clusters that overlaps with their 5’boundary. Current genome annotation of *S. cerevisiae* (R64-2-1) does not include several other types of non-coding genes transcribed by pol-II, such as “stable unannotated transcripts” (SUTs), “cryptic unstable transcripts” (CUTs) (Neil et al. 2009; Xu et al. 2009) and Xrn1-sensitive antisense regulatory non-coding RNA (XUT) (van Dijk et al. 2011). It is possible that some tag clusters belong to these non-coding transcripts, we included 847 SUTs, 925 CUTs and 1658 XUTs for cluster assignment. Gene Ontology (GO) term enrichment analysis was carried out by Go-TermFinder (https://go.princeton.edu/cgi-bin/GOTermFinder). GO Terms with P-values < 0.01 were classified as “significantly enriched”.

### Core promoter shift and gene differential expression analyses

Using tag distribution data obtained from YPD as a control, we estimated if the distribution of the CAGE tags between the top two active core promoters of a gene changes in other growth conditions. We implemented Chi-square test to infer whether there is a significant shift in term of tag distribution. We identified differential expressed genes in all eight comparisons using DESeq2 (Love et al. 2014). In both promoter shift and differential gene expression analyses, *p*-value of Chi-square test was adjusted with BH method (Benjamini & Hochberg) account for the multiple comparisons issue. Significant core promoter shift and differential expressed gene was defined as adjusted *p*-value less than 0.05.

### Promoter Shape Score

We developed a new metric, Promoter Shape Score (PSS), to quantify the shape of a core promoter based on the distribution of CAGE tags within a core promoter and promoter width. The PSS was calculated using the following equation:

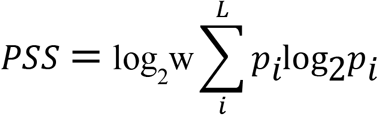

where *p* is the probability of observing a CTSS at base position *i* within a core promoter; *L* is the set of base positions that have normalized TSS density ≥ 0.1 TPM; and *w* is the promoter width, which was defined as the distance (in base pairs) between the 10^th^ and 90^th^ quantiles. This width marks the central part of the cluster that contains >= 80% of the CAGE signal.

### Sequence context analyses and *de novo* promoter motif discovery and

Dinucleotide frequencies were calculated with sequences extracted with bedtools nuc from [-1,+1] of TSS in *S. cerevisiae* genome (Quinlan 2014). Background dinucleotide frequencies were calculated with Perl script. To predict putative sequence motifs within or near the core promoters identified in this study, we performed *de novo* motif discovery with HOMER (http://homer.ucsd.edu/homer/). The sequences were retrieved from -100, +50 of the dominant TSS from each core promoter. The predicted motifs were compared with known motifs of transcription factor binding site datasets obtained from (MacIsaac et al. 2006) and Saccharomyces Cerevisiae Promoter Database (SCPD) (Zhu and Zhang 1999) using Tomtom module (Gupta et al. 2007) of the MEME Suite (Bailey et al. 2015).

### Data availability

The raw CAGE sequencing data have been deposited in the NCBI Sequence Read Archive (SRA) (accession number SRP155983). The quantitative maps of TSS and core promoter generated by this study can be visualized and downloaded from the YeasTSS database (http://www.yeastss.org).

## ACKNOWLEDGEMENTS

This study was supported by the start-up fund and Beaumont Award from Saint Louis University to ZL. We thank Dr. Yong Xue, and Dr. Hong Qin for valuable comments for improvement of this manuscript.

